# Cooperative nucleotide binding in Hsp90 and the underlying mechanisms

**DOI:** 10.1101/113191

**Authors:** Philipp Wortmann, Markus Goötz, Thorsten Hugel

## Abstract

The function of the molecular chaperone Hsp90 depends on large conformational changes, rearrangement of local motifs, as well as the binding and hydrolysis of ATP. The complexity of the Hsp90 system impedes the detailed investigation of their interplay using standard methods. By the application of three-color single molecule FRET to Hsp90 and a reporter nucleotide, we directly observe cooperativity between the two nucleotide binding pockets in the protein dimer. Through allocating the microscopic states and extracting their kinetics, we identify the mechanisms underlying the cooperativity. Surprisingly, nucleotide binding affects several state transitions, which demonstrates the complexity of cooperativity in protein systems. The co-chaperone Aha1, known to accelerate Hsp90's ATPase activity, adds another layer of complexity by affecting transitions in a nucleotide-dependent and -independent manner.

## Introduction

The molecular chaperone heat shock protein 90 (Hsp90) is the most abundant protein found in cells, accounting for up to 1−2 % of all cytosolic protein under physiological conditions^1^. It is highly conserved and essential in all eukaryotes^2^. A set of more than 20 co-chaperones is known to regulate the protein's function, building a complex network of transient interactions^1^. This Hsp90 machinery is essential for the correct folding of many cellular proteins as well as the maturation of kinases and steroid hormone receptors. It has thus become a popular drug target with several inhibitors developed and put into clinical trial for cancer therapy^3^. However, the way Hsp90 fulfills this job remains enigmatic.

Hsp90 is a homodimer consisting of three domains, the N-, the middle-and the C-terminal domain (NTD, MD, CTD). The NTD can bind and hydrolyze ATP. Hydrolysis in Hsp90s is very slow, e.g. about 1 ATP/min for the yeast homologue^4^. While stably dimerized at the CTD^5^, the relative orientation of the domains is dynamic and the dimer exhibits transitions between globally closed and largely open conformations^6–8^.

Yet, the interplay between nucleotide binding, hydrolysis and conformational changes of Hsp90 is poorly understood so far. ATP binding and hydrolysis weakly affect the conformational equilibrium between the globally open and closed conformations of Hsp90^9,10^. Only incubation with the non-hydrolysable ATP transition state analogue AMP-PNP leads to a stable closed structure of the otherwise predominately open Hsp90 dimer^7^.

Several bulk experiments and mutational studies have provided controversial results on the relationship between nucleotide binding, structural rearrangements and ATP hydrolysis^5,11–13^.

The standard procedure to access the interplay between two binding sites for a ligand (i.e. cooperativity) in a protein system is usually the evaluation of Hill-plots from bulk experiments, where the binding site occupation is measured as a function of ligand concentration. Cooperativity leads to a deviation from a slope of one in a log-log plot. However, this evaluation is limited by the simplifications of the underlying models^14^ and weak cooperativity is hardly detectable (Supplementary Results, Supplementary Fig. 1). But in the case of Hsp90, most interactions with clients, co-chaperones and nucleotides do not exhibit strong effects on its dynamics. Therefore, this procedure is not suitable for the characterization of any possible cooperative effects in Hsp90.

By applying three-color single molecule Förster Resonance Energy Transfer (smFRET)^15–17^ we are now able to resolve the microscopic states of the interaction between Hsp90 and nucleotides. Together with the kinetic description this enables us to detect effects that were previously hidden in the averaged ensemble data.

Our data reveals the first direct evidence of cooperativity between the two nucleotide binding pockets in the Hsp90 dimer. We can quantify the diverse effects on the microscopic state transitions of Hsp90 caused by bound nucleotide. This allows us to trace the mechanisms underlying the cooperativity. Moreover, we find that the co-chaperone Aha1 modulates the two nucleotide binding pockets and their interplay by an additional mechanism.

## Results

### The nucleotide binding pockets of Hsp90 act cooperatively

We study the effect of a native nucleotide on the *trans* nucleotide binding pocket with fluorescently labeled AMP-PNP as reporter nucleotide (subsequently denoted as AMP-PNP*). In order to follow the nucleotide binding state of Hsp90 while tracking the conformation of Hsp90 at the same time, we perform three-color smFRET measurements on labeled Hsp90 (Fig. 1a). The Hsp90 dimers are attached to the surface of a flow chamber and studied in the presence of 25 nM AMP-PNP* as detailed in^17^. This approach ensures that only one AMP-PNP* is bound at a time and that the second nucleotide binding site is accessible to the unlabeled nucleotide. The partial fluorescence (PF, the extension of the FRET efficiency for multi-color experiments) calculated from the fluorescence intensities is directly related to the spatial inter-dye distance by the Foörster radius R_0_. To get the full information on all three distances between the three dyes, alternating laser excitation is used.

**Figure 1:**
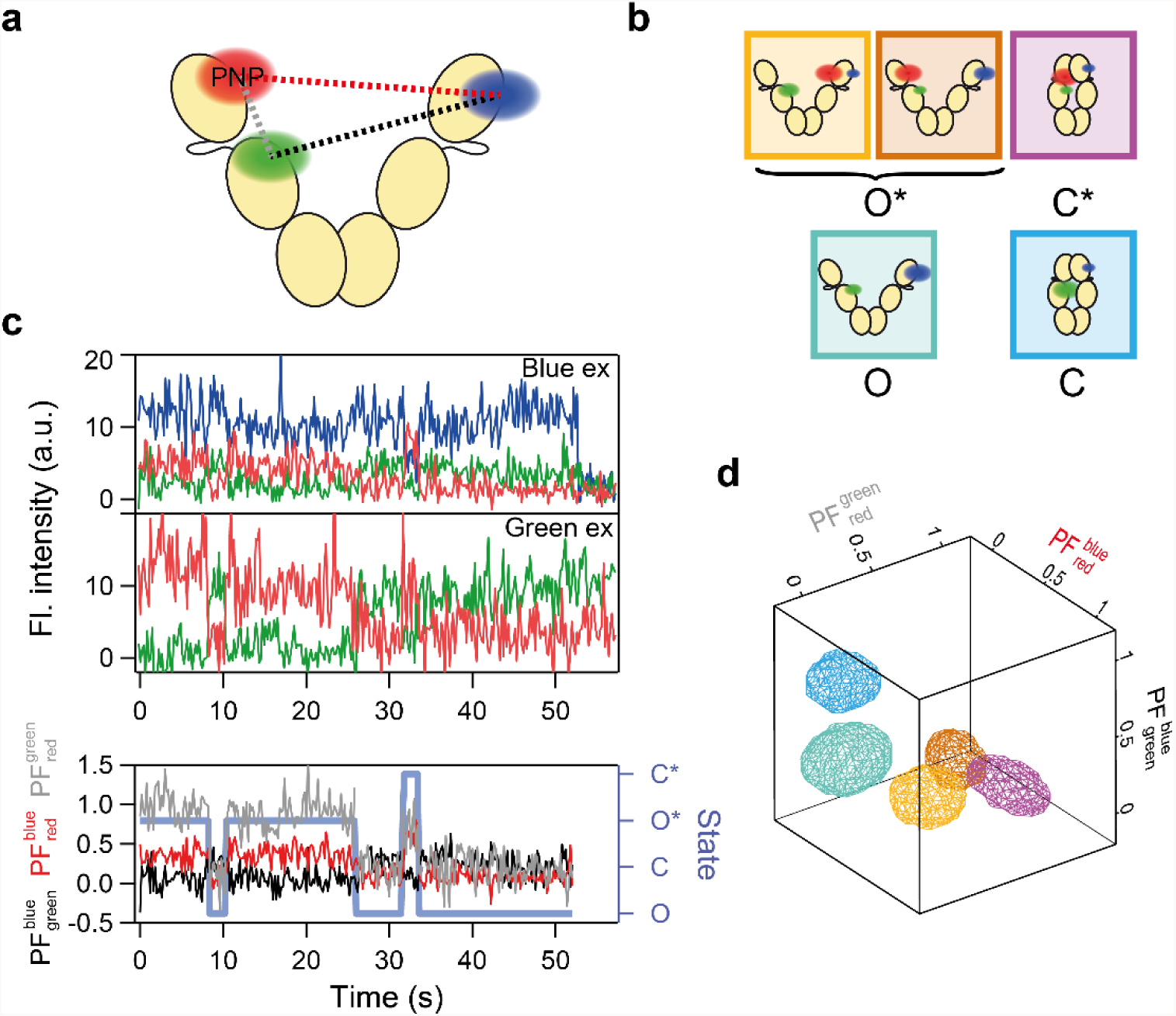
Data analysis and state allocation. (a) Pictogram of the studied system consisting of Hsp90 and the reporter nucleotide AMP-PNP* (PNP). (b) Pictograms of the distinguishable states and their identifier used in this work. The first two populations represent the same functional state O*. (c) Example fluorescence traces after the excitation of the blue (top) and the green (center) dye. The partial fluorescences (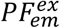, with excitation of the *ex* dye and emission of the *em* dye) calculated from the intensity traces are shown at the bottom. The state allocation is shown in light blue. (d) 3D representation of the Gaussians (isosurface at FWHM) fitted to the partial fluorescence data, which represent the five different populations. The color code is the same as in (b).

We are able to distinguish and allocate five different states to this system, as depicted in Figure 1b, of which four are functionally different. Namely, open nucleotide-free (O), open AMP-PNP* bound (O*), closed nucleotide-free (C) and closed AMP-PNP* bound (C*). The data (i.e. the fluorescence intensity traces; Fig. 1c) spans a three-dimensional space of the different PFs, which we subsequently use for state separation (Fig. 1d).

We collect traces from more than 400 single molecules that exhibited single AMP-PNP* binding under each experimental condition. A three-dimensional Hidden Markov Model (HMM) is used to allocate states in the individual smFRET time traces and extract a kinetic model from the data^17^.

With our single molecule approach we are able to measure the time for a single AMP-PNP* to stay bound to Hsp90 (Fig. 2a). Analyzing more than 800 dissociation events, we find that the average dwell time of AMP-PNP* on Hsp90 is 5.9 ± 0.3 s (Fig. 2b). If no cooperativity between the two binding pockets of Hsp90 existed, the addition of unlabeled nucleotide would not affect the average dwell time of the reporter already bound to Hsp90. Surprisingly, under conditions with additional, native ATP or AMP-PNP, the average dwell time is increased significantly by almost 50 % to 8.8 ± 0.3 s. Thus, the binding of a second nucleotide to the Hsp90 dimer decreases the apparent overall dissociation probability of the reporter nucleotide AMP-PNP* (averaged over all conformations of Hsp90), i.e. the two nucleotide binding pockets are not independent of each other. In other words, there exists cooperativity between the two binding pockets of Hsp90.

**Figure 2:**
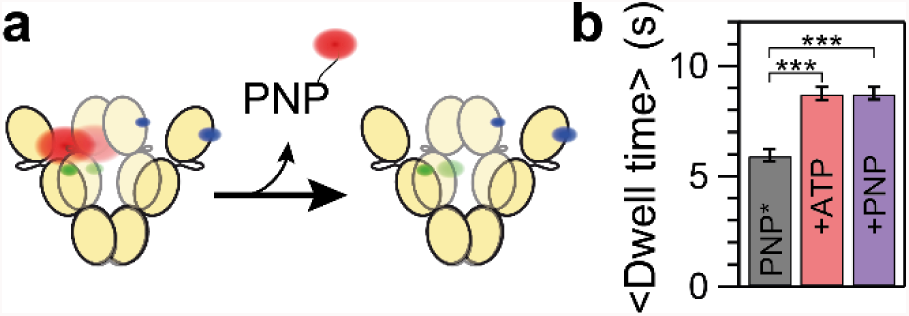
The average dwell time of the reporter nucleotide AMP-PNP* bound to Hsp90 is prolonged by additional nucleotide. (a) Pictogram of the observed dissociation of labeled AMP-PNP* (PNP) from the Hsp90 dimer. (b) Average dwell time of AMP-PNP* bound to Hsp90 without additional nucleotide (black) and with ATP (red) or AMP-PNP (purple) present. Error bars represent the standard deviations estimated from jackknife resampling. Differences between the dwell time distributions are significant with a p-value<0.001 (***).

As Hsp90 is a complex system with at least two distinct global conformations and several local motifs that could mediate this effect, a more detailed analysis is necessary in order to be able to identify its cause.

### Nucleotides have multiple effects on Hsp90

Figure 3a shows, that the ratio of Hsp90 with AMP-PNP* bound in the open and the closed state (O*/C*) is shifted towards the closed state in the presence of additional nucleotide. Hence, ATP and AMP-PNP have a similar effect on both the mean dwell time of AMP-PNP* and the population shift between O* and C*. However, the underlying mechanism of this effect remains unclear at this stage. The increase in the population of C* could be due to an increase in the rates leading to this state or due to a decrease in the rates depopulating this state. Accordingly, the increase in the average dwell time of bound AMP-PNP* may be caused by the shift from O* to C* (which has a slower dissociation rate of PNP*) or by a decrease in the dissociation rate from O* - or by a combination of both. Only a complete kinetic description can provide information about the actual mechanism mediating cooperativity in Hsp90. For such a multi-state system, this is currently only possible by a single molecule approach.

**Figure 3:**
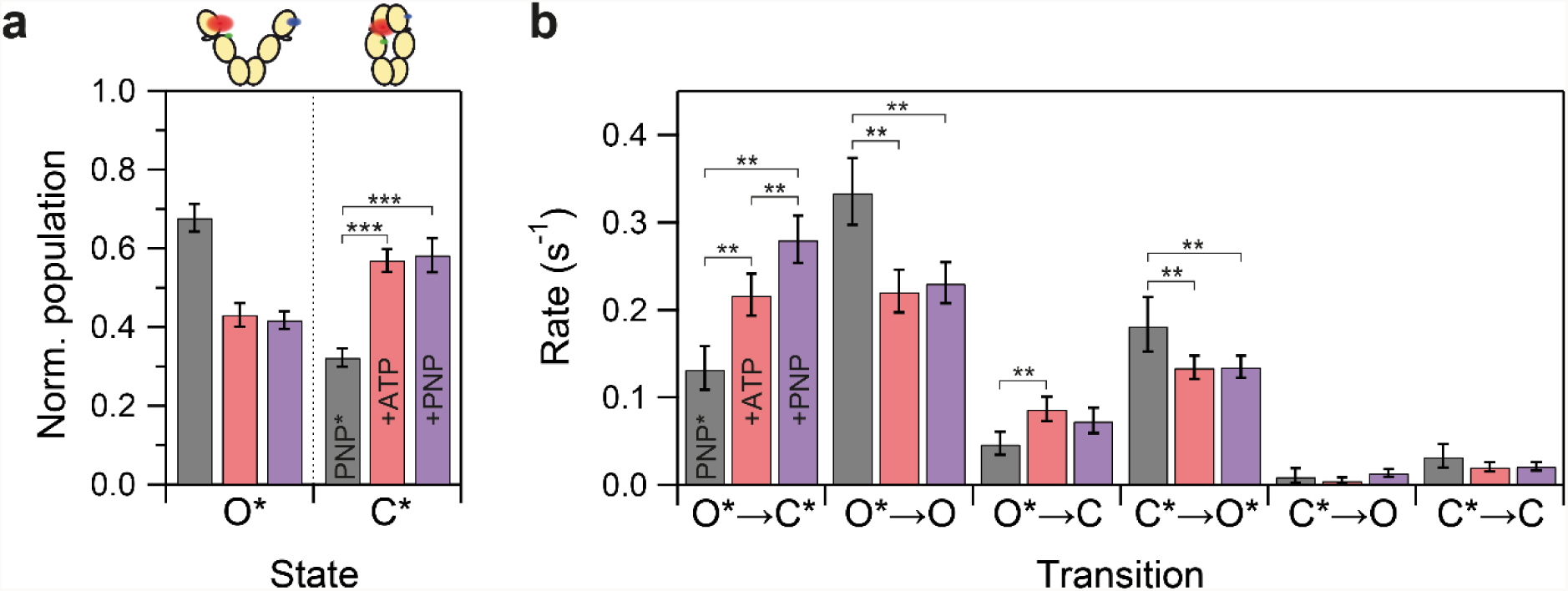
The effects of nucleotide on Hsp90's conformation and its state transitions. (a) Populations of open and closed conformation for Hsp90 bound to AMP-PNP* (normalized to unity) without additional nucleotide (black) and in presence ATP (red) or AMP-PNP (purple). Error bars represent the standard deviation within ten subsets, each comprising 75 % of the full dataset. The addition of nucleotide results in a significant population shift with a p-value<0.001 (***). (b) Transition rates for Hsp90 bound to labeled AMP-PNP* in dependence of additional nucleotide. Error bars represent the 99 % confidence interval (CI), differences with p<0.01 (CIs do not overlap) are highlighted (**).

Our three-color smFRET data enables us to simultaneously resolve the microscopic rates for conformational changes and AMP-PNP* unbinding for Hsp90. As shown in Figure 3b, additional nucleotide affects the nucleotide dissociation and the conformational transitions differently. On the one hand, additional nucleotide decreases the rates for AMP-PNP* dissociation from both conformational states of Hsp90 (O*→O and to a lesser extent C*→C). On the other hand, the equilibrium between O* and C* is shifted by both, increasing the rate for closing (O*→C*) and decreasing the rate for opening (C*→O*). Since the bleach rates in all data sets are similar, this effect is not introduced into our data by different bleaching times (Supplementary Fig. 2).

Thus, cooperativity is not caused by the nucleotide affecting one single rate, but by the combined effects of the nucleotide on four rates (indicated by ** in Fig. 3b) in the studied system.

### Aha1 and nucleotide binding affect Hsp90 by independent and interfering mechanisms

The ATPase activity of Hsp90 is known to be affected by regulatory co-chaperones. The strongest stimulating effect has been found for Aha1 so far, with a more than tenfold acceleration of the ATPase activity^18^. Since nucleotide binding is a prerequisite for hydrolysis, the question arises whether Aha1 affects the cooperativity of this process.

Therefore, we test the effect of Aha1 on the cooperativity in nucleotide binding by the addition of 10 μM of wild-type Aha1 to our assay. In absence of additional ATP, Aha1 alone already increases the mean dwell time of the reporter nucleotide bound to Hsp90 from 5.9 ± 0.3 s to 7.7 ± 0.3 s (Fig. 4a). Surprisingly, the combination of Aha1 and ATP increases the dwell time of AMP-PNP* (7.0 ± 0.3 s) to a lesser degree than either Aha1 or ATP do. Thus, the effects of ATP and Aha1 are not additive.

**Figure 4:**
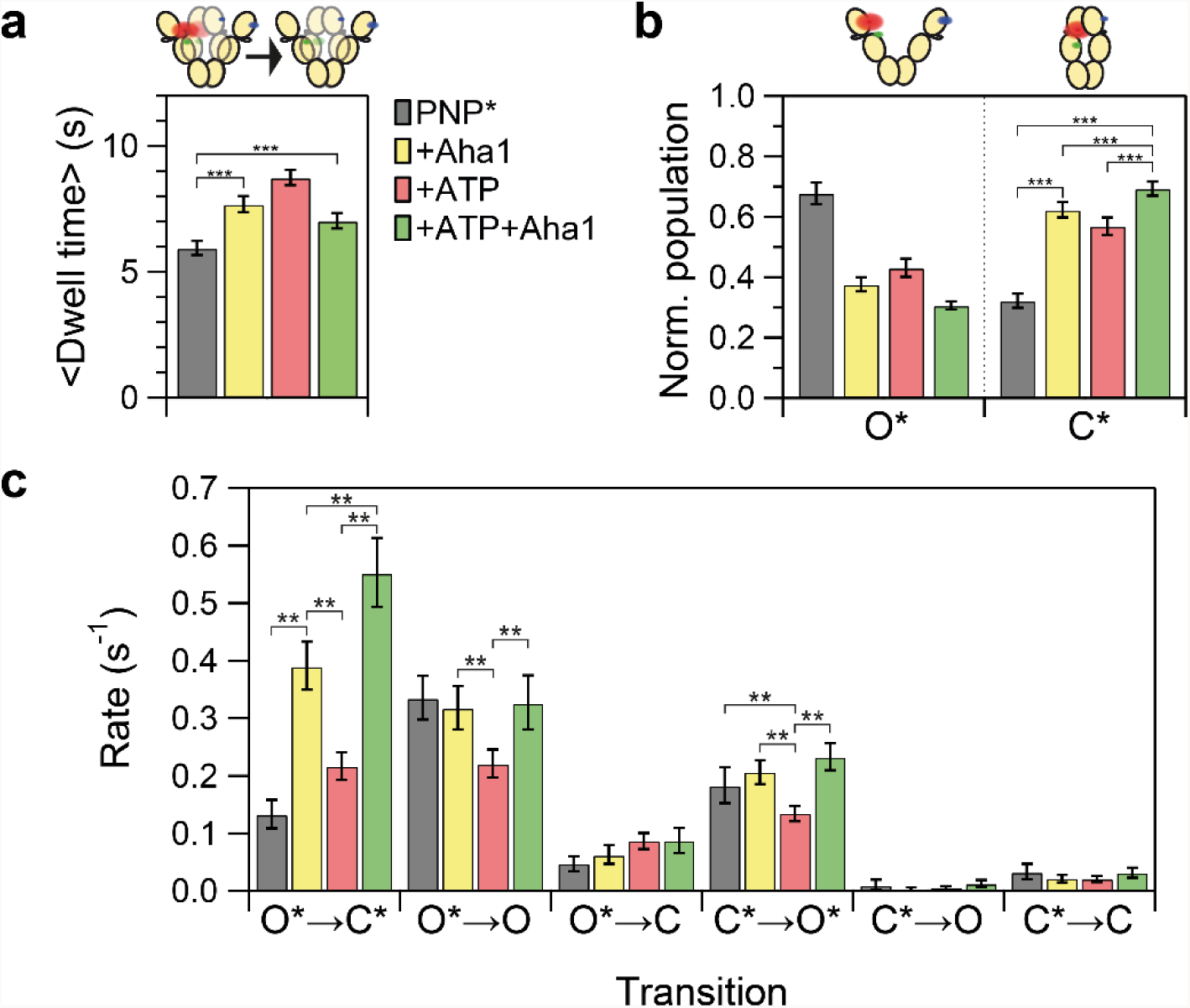
The effects of Aha1 (yellow), ATP (red) and Aha1 combined with ATP (green) on Hsp90 and AMP-PNP*. (a) The mean dwell time of AMP-PNP* on Hsp90 is significantly (p < 0.001, ***) increased by Aha1. The effect observed for Aha1+ATP is smaller than for Aha1 or ATP alone. Error bars are calculated as in Fig. 2. (b) The effects on the normalized population of open and closed conformation for Hsp90 bound to AMP-PNP*. Error bars are calculated as in Fig. 3a. (c) The transition rates and the effects of nucleotide, co-chaperone and their combination. Errors bars represent the 99 % CI, differences with p<0.01 are highlighted (**).

Figure 4b shows, how the O*/C* ratio of nucleotide bound Hsp90 is shifted towards the C* state by the addition of Aha1. In contrast to our findings for the dwell times, the combination of ATP and Aha1 leads to a more pronounced effect on the O*/C* ratio than each of them separately. Hence, Aha1 and ATP affect independent processes of Hsp90 and these processes interfere with each other.

The HMM analysis of the three-color smFRET data (Fig. 4c) provides a more detailed understanding of the microscopic mechanisms causing our observations. Aha1 alone accelerates the transition from open, AMP-PNP* bound Hsp90 (O*) to the closed state (C*), even stronger than ATP. In combination of Aha1 with ATP, this effect from both actually adds up. In contrast, the transitions from closed Hsp90 with AMP-PNP* bound to the open state as well as the dissociation of labeled nucleotide from open Hsp90 are increased to values similar to the ones observed without ATP and Aha1. Therefore, here Aha1 attenuates the additional effects of ATP.

## Discussion

The presented assay, based on the binding of the reporter nucleotide AMP-PNP* to one protomer of Hsp90, provides for the first time a detailed insight into the cooperation of the two nucleotide binding sites in the Hsp90 dimer. AMP-PNP* is not hydrolyzable and binds to the nucleotide binding pocket of Hsp90 with higher affinity than the unlabeled nucleotide (Supplementary Fig. 5 and 6). This makes it a highly useful reporter for an otherwise hardly accessible state of Hsp90, namely the state in which exactly one NTD adopts a conformation that presumably resembles the one that occurs during hydrolysis.

The observed cooperativity between the nucleotide binding sites explains several previously observed effects of point mutations in one of the nucleotide binding pockets of functional Hsp90 heterodimers.

That is, the effects of point mutations impairing nucleotide binding (D79N) or hydrolysis (E33A) on the ATPase rate vary significantly^5,10–12^. On the one hand, an Hsp90 heterodimer able to bind only one ATP exhibits an increased Michaelis-Menten constant K_M_^5,11^. Hence, in absence of a second ATP, the apparent nucleotide affinity of Hsp90 drops. The apparent nucleotide affinity in ensemble experiments always represents combined affinities, because Hsp90 exists in an equilibrium of at least two conformational states with the ability to bind ATP^22^. On the other hand, an Hsp90 heterodimer able to bind two ATPs, but with the ability to hydrolyze only one, displays a decreased K_M_ (i.e. an increased apparent nucleotide affinity)^11^. In conclusion, the presence of a non-hydrolyzable nucleotide increases the apparent nucleotide affinity, while its absence decreases it. Both findings are readily explained by our observation of cooperativity among the two nucleotide binding pockets. If one binding pocket is occupied by a non-hydrolyzable nucleotide, the addition of ATP and thus the occupation of the second binding pocket with ATP results in a prolonged binding of the nucleotide. According to our proposed model (Fig. 5a), this results in an increase in closed nucleotide-bound Hsp90. That is in turn the conformational prerequisite for hydrolysis, since a correlation between population of the closed conformation of Hsp90 and ATPase rate has been reported^10,23^. The cooperativity between the two pockets is an additional aspect of Hsp90's cross-monomer coordination, which happens before the actual hydrolysis of nucleotide^24^.

**Figure 5:**
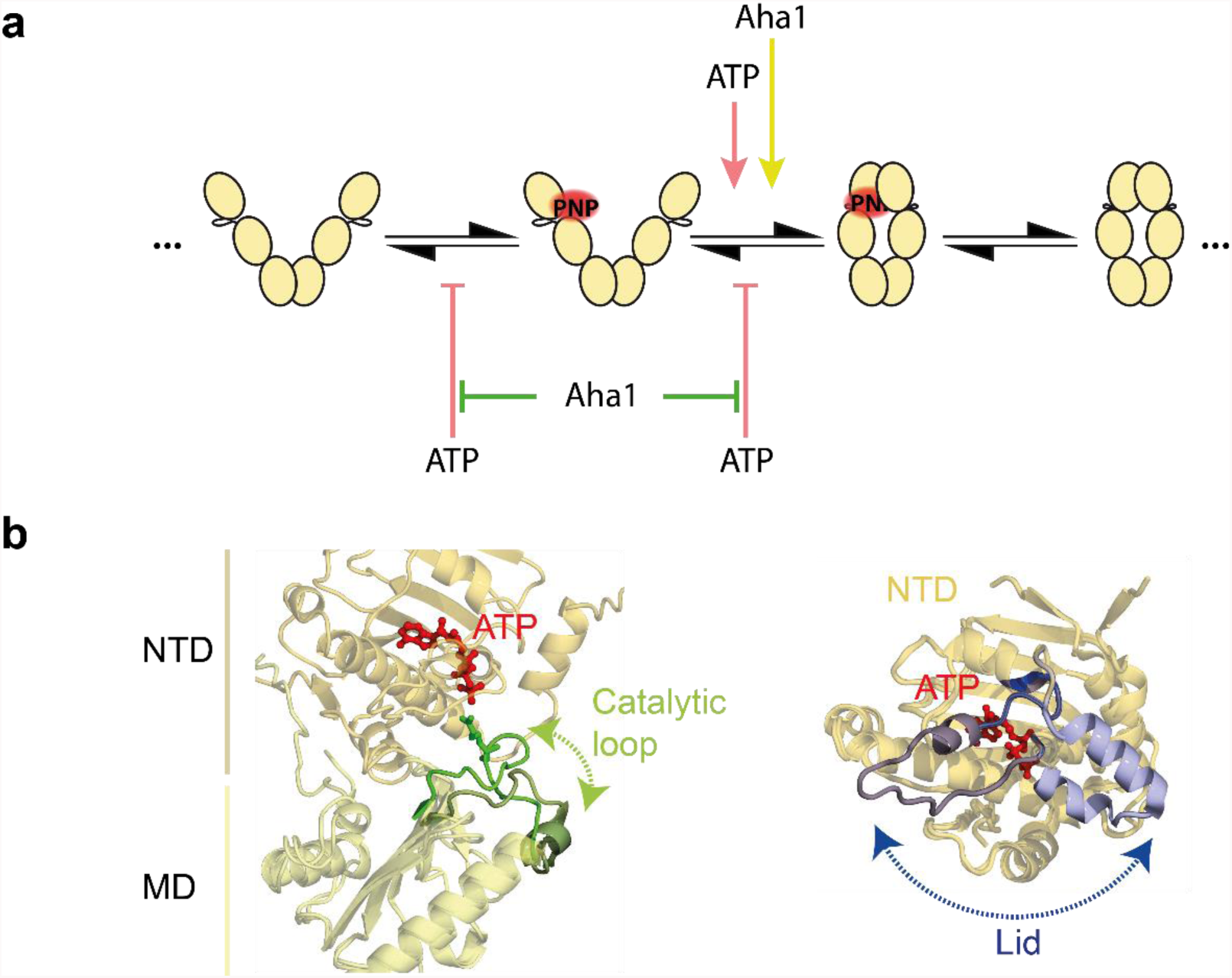
The effects of ATP and Aha1 on the state transitions of Hsp90 and their potential molecular origins. (a) A minimal model for the state transitions of Hsp90 in presence of AMP-PNP* and the effects of ATP and Aha1 on these transitions. Only the most frequent transitions are shown. ATP increases the closing rate of Hsp90 with AMP-PNP* bound, and decelerates the reverse reaction, as well as the dissociation of AMP-PNP*. Aha1 also accelerates the closing of AMP-PNP* bound Hsp90. In a combination of Aha1 and ATP, their effects on the closing of AMP-PNP* bound Hsp90 add up, while Aha1 prevents the decelerating effects of ATP. (b) Our observations could be caused by the depicted local rearrangements, which have been proposed to be affected by Aha1 and nucleotide binding. Left, reported rearrangement of the catalytic loop upon binding of Aha1 (green and dark green, superposition of the crystal structures PDB 2cg9^19^ and 1osv^20^). Right, reported conformation of the nucleotide lid in the AMP-PNP or the ADP bound crystal structures (dark and light blue, super position of the crystal structures PDB 2cg9 and 2wep^21^).

Our data also allows a deeper insight into the microscopic causes for the cooperativity. The addition of ATP accelerates the closing of AMP-PNP* bound Hsp90 (O*→C*), it decelerates the opening of AMP-PNP* bound Hsp90 (C*→O*) and the dissociation of AMP-PNP* from open Hsp90 (O*→O). The latter could be achieved either by a communication between the two nucleotide binding sites through the whole dimer^12,25^ or by a direct interaction between the N-domains. Direct steric contacts of the two NTDs in the open state are possible, because the open conformation of Hsp90 represents a flexible and highly dynamic ensemble^7^. Interestingly, a stable N-terminal closure of Hsp90 has been reported for saturating AMP-PNP conditions but not for ATP^6,9,10^. Here, we see that the presence of AMP-PNP in one binding pocket and the absence of nucleotide in the other is not sufficient for a more pronounced closed state of Hsp90. However, the presence of ATP in one binding pocket and AMP-PNP in the other results in a larger population of the closed state. Thus, binding of one AMP-PNP results in a conformational change that is the prerequisite for a more stable N-terminal dimerization. ATP has been proposed to affect the lid dynamics of Hsp90^26^, which can then facilitate the N-M arrangement necessary for the transition into the closed Hsp90 conformation. This molecular arrangement would explain the decreased opening rate (C*→O*) as well.

Furthermore, our data reflects the role of Aha1 as an accelerator of Hsp90's conformational rearrangements and ATPase activity^18,27^. We find that Aha1 accelerates the transition from the open nucleotide bound (O*) to the closed nucleotide bound state (C*) in absence of additional nucleotide. As only the forward reaction is affected (O*→C*) while the reverse is not, Aha1 is not acting in an enzyme-like fashion on this transition. Instead, Aha1 decreases not only the energy barrier of the O*→C* transition, but it also affects the equilibrium between these two states. This shift of AMP-PNP* bound Hsp90 towards the closed state in our experiments matches the previous finding of a stronger binding of Aha1 to the latter conformation^28,29^. Aha1 has been reported to modulate the catalytic loop of Hsp90 to facilitate the conformational transition from open to closed Hsp90^20^ (Fig. 5b). This is could be the microscopic cause for the effect we observe.

The addition of both, ATP and Aha1 to our assay allows further and novel insight into the distinct effects of nucleotide and co-chaperone on Hsp90's conformational dynamics. Their common and strongest effects, accelerating the closing of open, AMP-PNP* bound Hsp90 (O*→C*), add up. Because the concentration of Aha1 in our experiment is close to saturation^18,28,30^, we assume that the co-chaperone is already exerting its full activation potential. Nevertheless, the transition can be further accelerated by the addition of ATP. This suggests that ATP and Aha1 affect different structural rearrangements independently, both resulting in an accelerated closing of Hsp90. The decelerating effect of ATP on the opening of AMP-PNP* bound Hsp90 and dissociation of AMP-PNP* from open Hsp90 is abolished by Aha1. Thus, a structural arrangement leading to a tighter binding of nucleotide when both binding pockets are occupied is impeded by the co-chaperone. Therefore, on the one hand, ATP and Aha1 work to the same aim (namely closure of the dimer). On the other hand, they act antagonistic on two transitions (i.e. opening of dimer and dissociation of nucleotide). Thus, we conclude that at least two different structural motifs of Hsp90 are involved in their interaction. Apart from the aforementioned catalytic loop, the ATP lid might be influenced by both, ATP and Aha1^30^.

Our single molecule data and analysis enables us to conclude that the previously observed overall accelerating effect of Aha1 on Hsp90's conformational transitions^29,31^ is the result of at least two distinct and interfering modulations by the nucleotide and the co-chaperone. This demonstrates that simplifications assuming co-chaperones affect only single state transitions of Hsp90 do not reflect the complexity of this chaperone system. This possibly applies to many other catalytic systems with more than one binding site.

As a chaperone, Hsp90 interacts with folding intermediates of client proteins^1^. A recently published structure shows closed Hsp90 separating two domains of a kinase for correct folding and maturation in form of a molecular clamp^32^. In this context, we speculate that Aha1 together with ATP could increase the frequency of attempts of Hsp90 to separate the two domains of a client as an essential step in correct protein folding or maturation.

## Methods

All chemicals were purchased from Sigma-Aldrich if not stated otherwise.

### Protein expression, purification, labeling

Hsp90 from S. *cerevisiae* (UniProt ID P02829) is expressed in form of two single cysteine point mutants at the positions 61 and 385, with a cleavable N-terminal His-SUMO-tag and a C-terminal zipper ensuring the dimeric nature of the protein at picomolar concentrations. In the case of the 385C mutant, the construct contains an additional AviTag (Avidity LLC) at the extreme C-terminus for *in vivo* biotinylation^9^. Proteins were expressed and purified as described in^17^.

Hsp90 61C is labeled in 1× PBS pH 6.7 with Atto488-Maleiimide, Hsp90 385C with Atto550-Maleiimide (AttoTec GmbH), following the protocol provided in^17^. In case of Atto488, additional 0.5 mg/ml BSA are present.

Aha1 from S. *cerevisiae* (UniProt ID Q12449) is expressed and purified as wild-type construct with a cleavable N-terminal His-SUMO tag from pET28a. Transformed *E. coli* BL21Star (DE3) (Thermo Fischer Scientific) are grown at 37 °C in TB_Kan_ to OD_600_ of 0.6 and induced for 4 h. Cells are harvested by centrifugation, washed with phosphate buffered saline and lysed in a Cell Disrupter (Constant Systems) at 1.6 kbar. After filtration, the cell debris is applied to a 5 mL HisTrap HP (GE Healthcare) and eluted by a linear gradient from 0 to 500 mM imidazole in 100 mM sodium phosphate, 300 mM NaCl pH 8.0 at 8 °C. Protein-containing fractions are pooled and dialyzed into the imidazole-free buffer overnight in the presence of 1/100 mol/mol Senp protease to cleave the tag. The solution is again applied to the HisTrap column and the flow-through is collected and diluted 1:3 in 40 mM MES, pH 6, 40 mM NaCl and subsequently applied to a HiTrapSP 5 mL column (GE Healthcare). The protein is eluted with a linear gradient to 1 M NaCl. Protein is reduced with 1 mM DTT and concentrated. Finally, it is applied to a HiLoad 16/600 Superdex200 (GE Healthcare) and eluted with 40 mM Hepes, 200 mM KCl pH 7.5. Sample purity is checked by SDS-PAGE.

### ATPase assay

ATPase activity is determined with a regenerating ATPase assay. ATPase rates are measured at 37 °C with 2 mM ATP in 40 mM Hepes, 150 mM KCl, 10 mM MgCl_2_, pH 7.5. ATPase background is detected by specific inhibition of Hsp90 with radicicol and subtracted.

### Fluorescence anisotropy

Fluorescence anisotropy of labeled nucleotide is measured on a fluorescence spectrometer (Fluoromax-4, Horiba) with a temperature control unit (TC425, Quantum Northwest) at 25 °C. The label Atto647N is excited at 630 nm, fluorescence is detected at 660 nm with excitation and detection slits set to a width of 2 nm. The G-factor is determined prior to titration of protein and kept constant for the correction of all titration data points.

### Three-color smFRET measurements

Heterodimers of Hsp9061C-Atto488 and Hsp90_Biotin_385C-Atto550 are formed by the incubation of an overall 1 μM 5:1 mix for 20 minutes at 47 °C. Aggregates are subsequently removed by extensive centrifugation (> 30 minutes 17,000 xg).

Heterodimers are flushed into the custom built flow chamber depicted in^17^ at an appropriate concentration (approx. 10 pM) to reach sufficient surface density. Unbound protein is flushed out with buffer. 25 nM AMP-PNP-647N (AMP-γ-(6-Aminohexyl)-PNP-Atto647N, Jena Bioscience), here called AMP-PNP*, and 250 μM unlabeled nucleotide and/or 10 μM co-chaperone are flushed into the chamber. This step is repeated after 5 minutes of incubation again. All experiments are conducted in 40 mM Hepes, 150 mM KCl, 10 mM MgCl_2_, pH 7.5 and 0.5 mg/ml BSA.

smFRET traces are recorded by alternating laser excitation with a multi-color smFRET prism-type TIRF setup, as detailed in^17^. Shortly, Atto488 and Atto550 are excited with a blue (Cobolt Blues 50 mW, Cobolt) and a green (Compass 215M 75 mW, Coherent Inc.) laser at 473 and 532 nm, and fluorescence in the respective blue, green and red (for Atto647N) emission channels is detected by two EMCCD cameras (iXon Ultra 897, Andor Technology Ltd). Shutters, AOTF and cameras are synchronized by a digital I/O card (PCIe-6535, National Instruments). The overall time resolution of our experiments is 200 ms per excitation cycle, with each excitation lasting 70 ms (+30 ms read-out).

### Relative State population

We calculate the partial fluorescence 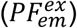 from the five intensity traces analogous to the FRET efficiency of a two-color experiment^17^. E.g. the partial fluorescence of AMP-PNP* (identifier “red”) after excitation of Atto488 (“blue”) is given by:

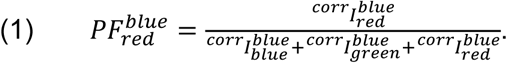

The relative state populations are determined from 2D projections of the PF histogram on the 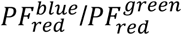 and 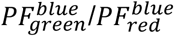 planes. Populations are selected by drawing free-hand polygons that represent the location of each state. This procedure was repeated three times to minimize the effect of manual selection.

### Ensemble HMM

The 3D histogram of the observed PFs reveals five different populations that are fitted by the sum of five 3D Gaussians with means and covariance matrix as fitting parameters. A global HMM was optimized for each data set with the emission probabilities for each state fixed to the 3D Gaussians determined by the fit.

The resulting transition matrix is multiplied with the frame rate to yield the rate constants of the transitions. Rates from functional equal states (the both O* states) were added for clarity.

Dwell times of the AMP-PNP* bound fraction are extracted from the Viterbi path. Dwells without recorded start and/or end where also included.

All data analysis was done with custom written scripts in Igor Pro (Wavemetrics). The code is available upon request.

### Error estimates

Dwell time distributions follow exponential functions. The mean of random samples from a dwell time distribution follows approximately a normal distribution for a large number of samples (central limit theorem). The variance of a normal distributed quantity that has been sampled n times can then be estimated by the jackknife-1 method, i.e. calculation from all possible combinations of n−1 dwells. It is then estimated by ^33^:

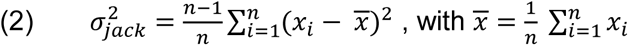

The error of the population size was estimated by the standard deviation of ten random subsets containing 75 % of the frames in the data set.

### Statistical tests

The dwell time distributions were post-hoc tested pairwise for significant differences by a one-sided Wilcoxon-Mann-Whitney two-sample rank test as implemented in Igor Pro (Wavemetrics). Test results are given in Supplementary Results, Supplementary Table 3.

The determined populations were tested for normal distribution by a Shapiro-Wilk test (as implemented in Igor Pro) on the subset distributions, results are given in Supplementary Results, Supplementary Table 2. The populations from different subsets were tested for significant differences post-hoc by an unpaired pairwise t-tests on the C* population. Results are given in Supplementary Results, Supplementary Table 1.

## Acknowledgments

We thank Dieter Hauschke and Jens Timmer for advice on data statistics. We thank Bizan Balzer, Björn Hellenkamp, Markus Jahn, Sonja Schmid, Katarzyna Tych and Matthias Rief for helpful discussions. This work is funded by the European Research Council through ERC grant agreement no. 681891.

## Author contributions

T.H., P.W. and M.G. designed research. P.W. and M.G. performed experiments and analyzed the data. P.W., M.G. and T.H. interpreted the data and wrote the manuscript.

## References

1. Taipale, M., Jarosz, D. F. & Lindquist, S. HSP90 at the hub of protein homeostasis: emerging mechanistic insights. Nature reviews. Molecular cell biology 11, 515–528 (2010).

2. Eckl, J. M. & Richter, K. Functions of the Hsp90 chaperone system: lifting client proteins to new heights. International Journal of Biochemistry and Molecular Biology 4, 157–165 (2013).

3. Jhaveri, K. et al. Heat shock protein 90 inhibitors in the treatment of cancer: current status and future directions. Expert opinion on investigational drugs 23, 611–628 (2014).

4. Panaretou, B. et al. ATP binding and hydrolysis are essential to the function of the Hsp90 molecular chaperone in vivo. The EMBO journal 17, 4829–4836 (1998).

5. Richter, K., Muschler, P., Hainzl, O. & Buchner, J. Coordinated ATP hydrolysis by the Hsp90 dimer. The Journal of biological chemistry 276, 33689–33696 (2001).

6. Mickler, M., Hessling, M., Ratzke, C., Buchner, J. & Hugel, T. The large conformational changes of Hsp90 are only weakly coupled to ATP hydrolysis. Nature structural & molecular biology 16, 281–286 (2009).

7. Hellenkamp, B., Wortmann, P., Kandzia, F., Zacharias, M. & Hugel, T. Multidomain structure and correlated dynamics determined by self-consistent FRET networks. Nature methods 14, 174–180 (2017).

8. Krukenberg, K. A., Forster, F., Rice, L. M., Sali, A. & Agard, D. A. Multiple conformations of E. coli Hsp90 in solution: insights into the conformational dynamics of Hsp90. Structure (London, England: 1993) 16, 755–765 (2008).

9. Schmid, S., Goötz, M. & Hugel, T. Single-Molecule Analysis beyond Dwell Times: Demonstration and Assessment in and out of Equilibrium. Biophysical journal 111, 13753–1384 (2016).

10. Zierer, B. K. et al. Importance of cycle timing for the function of the molecular chaperone Hsp90. Nature structural & molecular biology 23, 1020–1028 (2016).

11. Mishra, P. & Bolon, D. N. A. Designed Hsp90 heterodimers reveal an asymmetric ATPase-driven mechanism in vivo. Molecular cell 53, 344–350 (2014).

12. Obermann, W. M., Sondermann, H., Russo, A. A., Pavletich, N. P. & Hartl, F. U. In Vivo Function of Hsp90 Is Dependent on ATP Binding and ATP Hydrolysis. J Cell Biol 143, 901–910 (1998).

13. McLaughlin, S. H., Ventouras, L.-A., Lobbezoo, B. & Jackson, S. E. Independent ATPase activity of Hsp90 subunits creates a flexible assembly platform. Journal of molecular biology 344, 813–826 (2004).

14. Stefan, M. I. & Le Novere, N. Cooperative binding. PLoS computational biology 9, e1003106 (2013).

15. Person, B., Stein, I. H., Steinhauer, C., Vogelsang, J. & Tinnefeld, P. Correlated movement and bending of nucleic acid structures visualized by multicolor single-molecule spectroscopy. Chemphyschem : a European journal of chemical physics and physical chemistry 10, 1455–1460 (2009).

16. Lee, N. K. et al. Three-color alternating-laser excitation of single molecules: monitoring multiple interactions and distances. Biophysical journal 92, 303–312 (2007).

17. Goötz, M., Wortmann, P., Schmid, S. & Hugel, T. A Multicolor Single-Molecule FRET Approach to Study Protein Dynamics and Interactions Simultaneously. Methods in enzymology 581, 487–516 (2016).

18. Panaretou, B. et al. Activation of the ATPase activity of hsp90 by the stress-regulated cochaperone aha1. Molecular cell 10, 1307–1318 (2002).

19. Ali, M. M. U. et al. Crystal structure of an Hsp90-nucleotide-p23/Sba1 closed chaperone complex. Nature 440, 1013–1017 (2006).

20. Meyer, P. et al. Structural basis for recruitment of the ATPase activator Aha1 to the Hsp90 chaperone machinery. The EMBO journal 23, 511–519 (2004).

21. Prodromou, C. et al. Structural basis of the radicicol resistance displayed by a fungal hsp90. ACS chemical biology 4, 289–297 (2009).

22. Ratzke, C., Berkemeier, F. & Hugel, T. Heat shock protein 90's mechanochemical cycle is dominated by thermal fluctuations. Proceedings of the National Academy of Sciences of the United States of America 109, 161–166 (2012).

23. Halpin, J. C., Huang, B., Sun, M. & Street, T. O. Crowding Activates Heat Shock Protein 90. The Journal of biological chemistry 291, 6447–6455 (2016).

24. Cunningham, C. N., Krukenberg, K. A. & Agard, D. A. Intra-and intermonomer interactions are required to synergistically facilitate ATP hydrolysis in Hsp90. The Journal of biological chemistry 283, 21170–21178 (2008).

25. Morra, G., Verkhivker, G. & Colombo, G. Modeling signal propagation mechanisms and ligand-based conformational dynamics of the Hsp90 molecular chaperone full-length dimer. PLoS computational biology 5, e1000323 (2009).

26. Prodromou, C. et al. The ATPase cycle of Hsp90 drives a molecular 'clamp' via transient dimerization of the N-terminal domains. The EMBO journal 19, 4383–4392 (2000).

27. Prodromou, C. Mechanisms of Hsp90 regulation. Biochemical Journal 473, 2439–2452 (2016).

28. Retzlaff, M. et al. Asymmetric activation of the hsp90 dimer by its cochaperone aha1. Molecular cell 37, 344–354 (2010).

29. Li, J., Richter, K., Reinstein, J. & Buchner, J. Integration of the accelerator Aha1 in the Hsp90 co-chaperone cycle. Nature structural & molecular biology 20, 326–331 (2013).

30. Siligardi, G. et al. Co-chaperone regulation of conformational switching in the Hsp90 ATPase cycle. The Journal of biological chemistry 279, 51989–51998 (2004).

31. Schulze, A. et al. Cooperation of local motions in the Hsp90 molecular chaperone ATPase mechanism. Nature Chemical Biology 12, 628–635 (2016).

32. Verba, K. A. et al. Atomic structure of Hsp90-Cdc37-Cdk4 reveals that Hsp90 traps and stabilizes an unfolded kinase. Science (New York, N.Y.) 352, 1542–1547 (2016).

33. Shao, J. & Wu, C. F. J. A General Theory for Jackknife Variance Estimation. Ann. Statist. 17, 1176–1197 (1989).

